# A Computational Approach for Threonine Accumulation in *Escherichia coli* and Its Integraton as a Platform for Biosynthesis of High-Value Fine Chemicals

**DOI:** 10.1101/2025.10.29.685076

**Authors:** Tommaso Sassi, Gabriele Rebuzzini, Bas Teusink, Marco Vanoni, Luca Brambilla

## Abstract

Using genome-wide metabolic models and Flux Balance Analysis, we are devising an *Escherichia coli* strain to produce L-threonine through previously undescribed routes. *In silico* simulations and preliminary *in vivo* experiments show that diverting carbon flux away from serine metabolism may be beneficial to enhance threonine production. This system may be further explored not only to produce threonine itself, but also to develop an *Escherichia coli* platform capable of producing high value-added products which require threonine as a starting substrate. In order to do this we took advantage of the increased threonine production to sustain the production of the high value-added chemical 2,5-dimethylpyrazine (DMP) from glucose.

## Introduction

L-threonine is an amino acid of industrial interest both as a feedstock and as a precursor to other chemicals, such as 2,5-dimethylpyrazine, which is currently used as an additive for flavoring in the food industry and as a precursor for hypoglycemic drugs [1]. Despite this, today DMP is produced only via expensive, polluting chemical processes. In *Escherichia coli*, threonine biosynthesis originates from the aspartate family of amino acids. A major challenge in threonine overproduction is the complex regulation of carbon and nitrogen fluxes and the feedback inhibition of the key enzyme aspartokinase by threonine itself. Over the past two decades, numerous strategies have been developed to overcome these limitations, including the overexpression of biosynthetic genes, the attenuation of competing pathways, and the elimination of regulatory bottlenecks [2] [3] [4] [5] [6]. However, these approaches often rely on intensive trial and error experiments or on information derived from previously engineered producing strains. Our goal is to exploit the advances in genome-wide metabolic models (GEMs) computational analysis to rapidly identify novel strategies to increase L-threonine production in *Escherichia coli* by pure computational analysis of the model. Finally, we combined our newly discovered approach to the expression of oxidative enzymes to produce DMP directly from glucose, thus achieving the production of a high value-added chemical of industrial interest from an inexpensive substrate.

## Material and methods

### Strains and plasmids

For this study we used *Escherichia coli* BL21(DE3). The plasmids used are shown in Table 1. Cells were transformed via electroporation using a BioRad Gene Pulser II following a standard protocol for electrocompetent cells [7].

**Table 1:** Plasmids used in this study.

| Plasmid | Description | Source |
| --- | --- | --- |
| pETDuet[tdh/aao] | High-copy plasmid for the coexpression of threonine dehydrogenase and aminoacetone oxidase from *Streptococcus oligofermentans* | This study |
| pdCas9-bacteria | aTc inducible catalitically inactive Cas9 for gene silencing | [8] |
| pgRNA-THR | Constitutively induced sgRNA for CRISPRi targeting *serB*, *serC*, *pykA*, *tdcB*, *ltaE*, *tynA* | This study |

### Flux Balance Analysis

The iEC1372 W3110 genome-scale model of *Escherichia coli* was used as a template for Flux Balance Analysis (FBA) [9]. The model was expanded by adding the reactions of the DMP-producing pathway (aminoacetone oxidase, dehydration of 3,6-dihydro-2,5-dimethylpyrazine and corresponding metabolites). Simulations were performed using COBRAPy [10].

### Thermodynamic analysis

To determine which substrate is the best for DMP production, the eQuilibrator platform was used to perform Max-min Driving Force analysis on the pathways. As input, the flux distributions obtained with FBA were converted into SBtab and loaded in the eQuilibrator platform [11] [12] [13].

### Growth conditions and silencing

For gene silencing, we chose to use CRISPR interference technology (CRISPRi), which is based on a catalitically inactive Cas9 that blocks the transcription of a target gene without cutting the target sequence. This not only allows reversible silencing, but allows a gradual silencing of essential genes, avoiding a total loss of growth [8]. Cells were grown overnight in a Biostat B reactor filled with 1 L of M9 medium supplemented with 5 g/L glucose. The next day, 25 mL of broth was collected and mixed in a 250 mL flask with 25 mL of fresh M9 medium with aTc for dCas9 induction. After 24 h, supernatant samples were collected for quantification of threonine.

### Real-Time PCR

CRISPRi was validated via Real-Time PCR. mRNA was extracted from the cells using the Qiagen RNeasy® Mini Kit and quantified by NanoDrop One. Retrotrascription was performed using Bio-Rad iScript Advanced cDNA synthesis. Real-Time PCR was performed 5 times on each sample. The primers used were designed with Beacon Designer 8 and are shown in Table 2. For nornmalization control, we used the housekeeping gene *idnT*. Relative expression was calculated using the 2*^−^*^ΔΔ*Ct*^ method.

**Table 2:**
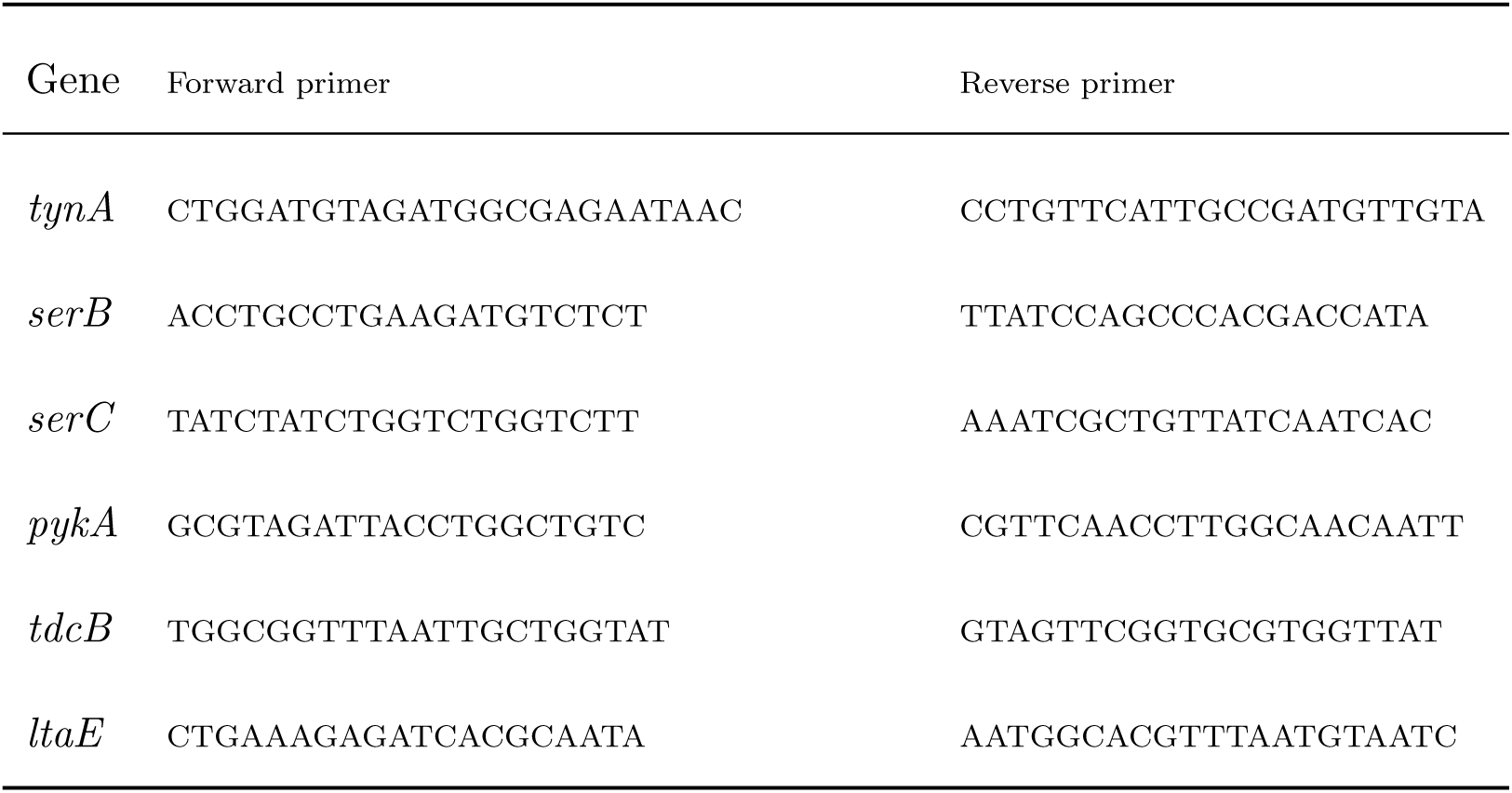
Primers used for Real-Time PCR.

### High Performance Liquid Chromatography

Threonine was quantified by HPCL using a modified version of the protocol by Handerson et al. [14]. Briefly, 5.5 *µ*L of sample were mixed with 5.5 *µ*L of derivatizing agent (5 mg/mL o-phtaldehyde and 5 mg/mL 3-mercaptopropionic acid in 0.2 M borate buffer pH 10.2). For detection a C18 kinetex EVO 5 *µ*m250 * 4.6 mm 100 Å column was used. A 40 mM sodium phosphate buffer (pH 7.8) and a 30:60:10 mix of acetonitrile, methanol and water were used as mobile phases. To measure DMP, a XTerra C18 4,6mm x 250 mm column was used in conjunction with a gradient protocol with trifluoroacetic acid 0.025% (phase A) and acetonitrile 100% (phase B). The gradient was programmed as such:

- Starting condition: phase A 100% - phase B 1%
- 0’ - 20’: phase A 92% - phase B 8%

Flux was kept at 1 mL/min and temperature at 25°C. Detection was performed with a UV lamp set at 260 nm.

### Liquid Chromatography Mass Spectrometry (LC-MS) Analysis

Threonine presence in the samples was also validated by mass spectrometry. Samples were injected into a Xevo G2-XS quadrupole time of flight mass spectrometer coupled to a Waters ACQUITY ultra-performance liquid chromatography (UPLC) H-Class system through an ESI source (Waters Corporation, Milford, MA, USA). The instrument was operated in positive ion current in sensitivity mode. The capillary voltage was maintained at 1 kV with the source temperature held constant at 140 °C. A Supel™ Carbon LC, 10 cm x 2.1 mm I.D., 2.7 *µ*m (MERCK) was used for analyte separation. The flow rate was set to 400 µL/min with a column temperature of 40 °C. A gradient of solvent A (99.9% LC-MS-grade water with 0.1% formic acid) and solvent B (LC-MS-grade 99.9% acetonitrile with 0.1% formic acid) was applied with a total run time of 13 min as follows: 0–3 min at 100% A; 3.5-7.5 min 100% B; 7.5-13 min 100% A.

## Results and discussion

### Identifying the best substrate for DMP production

In considering the synthesis of 2,5-dimethylpyrazine, the crucial starting point is L-2-aminoacetoacetate. The pathway is shown in Figure 1.

**Figure 1:**
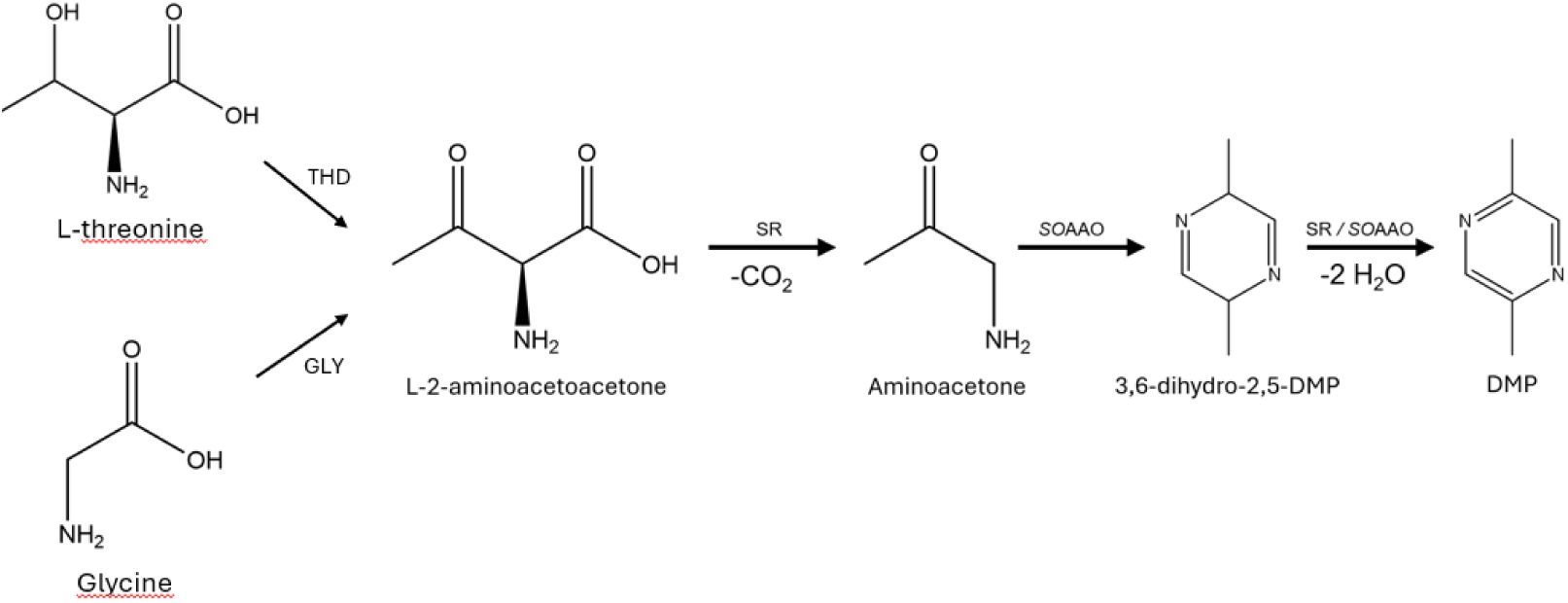
DMP production from L-2-aminoacetoacetone. THD = threonine dehydrogenase; GLY = glycine acetyltransferase; SOAAO = aminoacetone oxidase; SR = spontaneous reaction.

Despite in literature DMP production has been achieved from threonine via the enzyme threonine dehydrogenase, which oxidizes threonine to L-2-aminoacetoacetate, production from glycine is technically possible via glycine acetyltransferase [1]. For this reason, we applied a computational approach to assess which amino acid is the substrate that can ensure high DMP flux. The strategy followed two steps: 1) use Flux Balance Analysis (FBA) to define the flux distribution when L-2-aminoacetoacetate is produced from either glycine or threonine; 2) finding which flux distribution is actually optimal *in vivo*. We performed FBA on the iEC1372 W3110 genome-scale model of *Escherichia coli* by constraining the model to optimize production of DMP. By doing that, we force the model to produce at least the same amount of L-2-aminoacetoacetate, because all DMP eventually comes from this metabolite. We performed two different FBAs where DMP production was forced from either one of the two amino acids. After that, the two fluxes distributions were used as input for Max-min Driving Force (MDF) analysis on the eQuilibrator platform. The MDF value for a pathway is the smallest*−*Δ*_r_G^′^*obtained by any pathway reaction once all metabolite concentrations are chosen to make all pathway reactions as favorable as possible. Therefore, when comparing two pathways, the one with the highest MDF value will be the most favored thermodynamically. The cumulative *−*Δ*_r_G^′^* and MDF values of the two pathways are shown in Table 3.

**Table 3:** Thermodynamic values obtained by eQuilibrator. The threonine dehydrogenase route is the clear favorite.

| Pathway | Cumulative $\Delta_r G'$ | MDF |
| --- | --- | --- |
| Threonine dehydrogenase | -98.4 kJ/mol | -0.21 kJ/mol |
| Glycine acetyltransferase | 2.7 kJ/mol | -4.29 kJ/mol |

Because the glycine acetyltransferase route has the worst thermodynamic values of the two, we discarded this alternative and chose to investigate DMP production exclusively from threonine.

### Flux Balance Analysis for increased threonine production

After threonine was identified as the best substrate, we proceeded to find new ways to increase its production in *Escherichia coli*, since production of DMP needs to be sustained by a good flux of threonine biosynthesis. We performed two other FBAs, comparing the flux distributions obtained by maximization of biomass production and increased threonine production. With this comparison, we identified the fluxes that go to 0 when threonine production is maximized. We focused on these fluxes because fluxes that go to 0 in the simulations equate to fluxes that need to be silenced *in vivo*, which is straightforward to do. These fluxes are reported in Table 4.

**Table 4:**
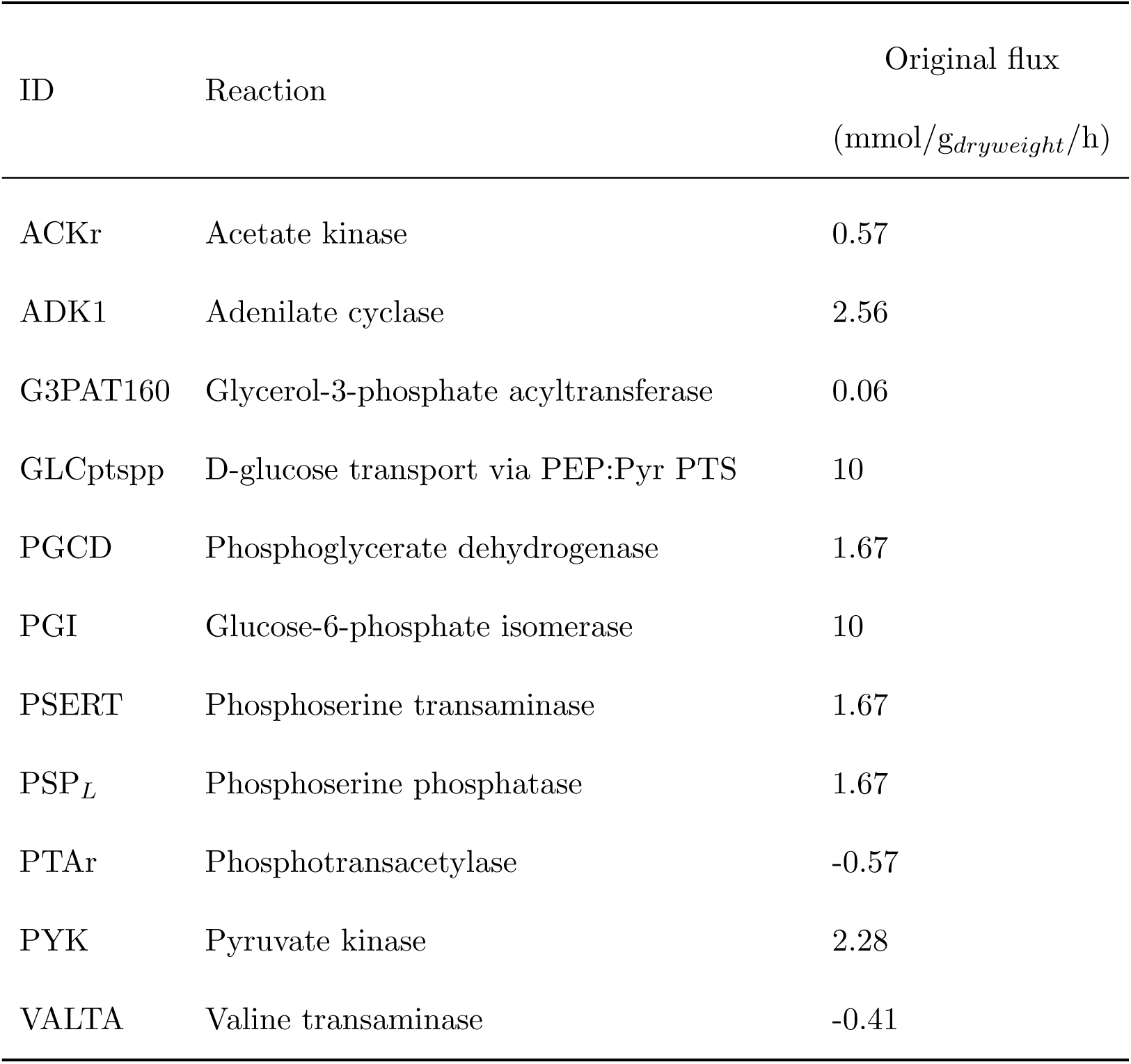
Reactions whose flux goes to 0 when threonine is increased. In the first column the model-specific ID of each reaction is reported.

| ID | Reaction | Original flux |
| --- | --- | --- |
|  |  | (mmol/g<sub>dryweight</sub>/h) |
| ACKr | Acetate kinase | 0.57 |
| ADK1 | Adenilate cyclase | 2.56 |
| G3PAT160 | Glycerol-3-phosphate acyltransferase | 0.06 |
| GLCptspp | D-glucose transport via PEP:Pyr PTS | 10 |
| PGCD | Phosphoglycerate dehydrogenase | 1.67 |
| PGI | Glucose-6-phosphate isomerase | 10 |
| PSERT | Phosphoserine transaminase | 1.67 |
| PSP<sub>L</sub> | Phosphoserine phosphatase | 1.67 |
| PTAr | Phosphotransacetylase | -0.57 |
| PYK | Pyruvate kinase | 2.28 |
| VALTA | Valine transaminase | -0.41 |

By manually simulating the deletion of each gene to evaluate how much the threonine production changes, we selected phosphoserine transaminase, phosphoserine phosphatase and pyruvate kinase as the best candidate for gene silencing, since they were responsible for the largest increase in threonine flux.

### Gene silencing and threonine production

For gene silencing in addition to *serB*, *serC* and *pykA* we decided to also target *ltaE*, *tdcB* and *tynA*. The first two genes codify threonine aldolase and threonine dehydratase, both responsible for consuming threonine, while *tynA* codifies for a primary amine oxidase which cataliyes the oxidation of aminoacetone to methylglyoxal, subtracting it from DMP synthesis. Overall, these three genes are detrimental to our goal. CRISPRi was our strategy of choice because it allows for a gradual, reversible and not total silencing, which is particularly suited for this case since because phosphoserine transaminase and phosphoserine phosphatase are both essential genes [15]. The results of real-time PCR and threonine levels are shown in Figure 2 and 3.

**Figure 2:**
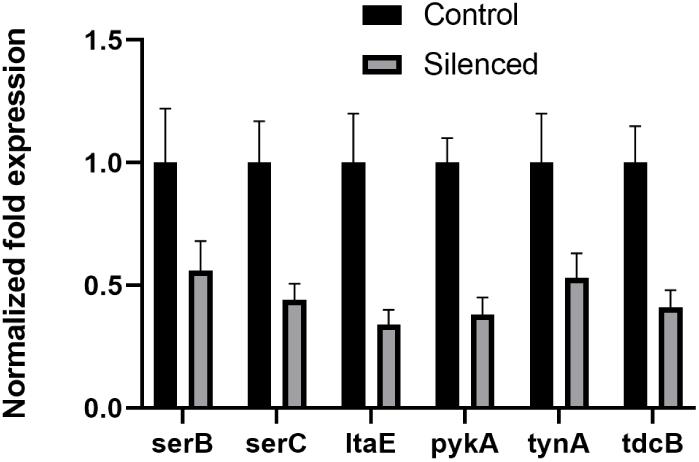
mRNA levels of the targeted genes after CRISPRi.

**Figure 3:**
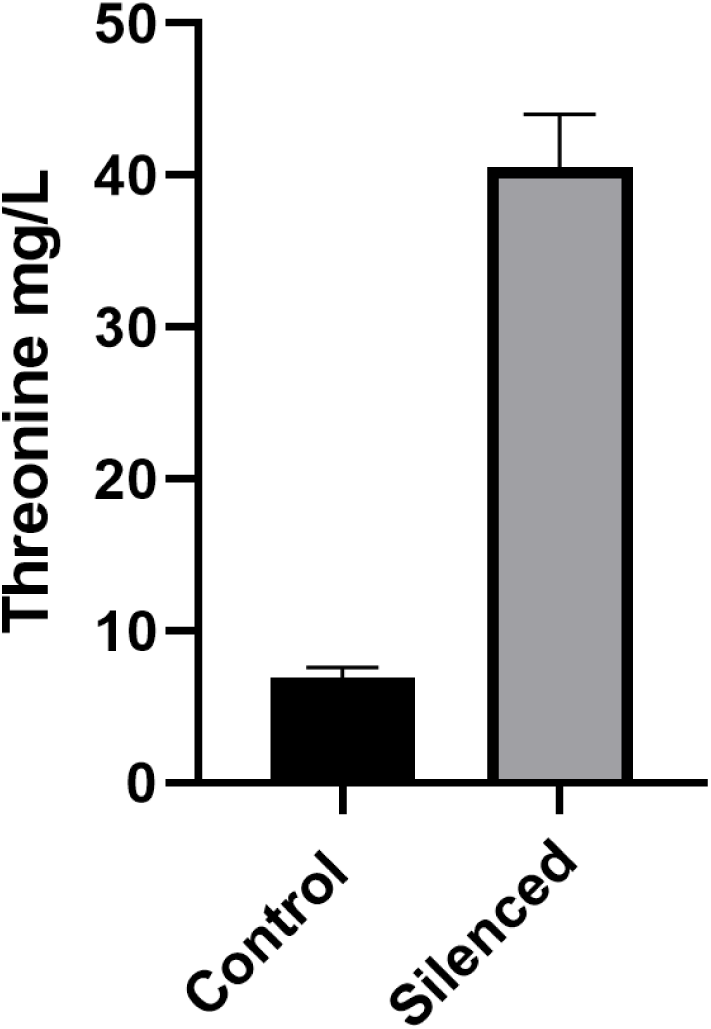
Extracellular threonine accumulation in flask after 24 h.

RT-PCR data shows a partial silencing of the genes, which is expected with CRISPRi, while HPLC analysis shows a 5.6-fold increase of extracellular threonine compared to the control. Threonine concentration was confirmed by MS analysis. Other works used silico approaches to optimize strains that were already engineered for threonine production [4], while we described a previously unknown strategy devised purely by analysis of the genome-scale metabolic model of *Escherichia coli*. The increase of threonine upon downregulation of serine biosynthesis can be explained by the overall competition of these two amino acids for carbon: serine biosynthesis starts from 3-phoshpoglycerate, while threonine biosynthesis starts from aspartate. Limiting the biosynthesis of serine can then leave more carbon available for aspartate, and therefore threonine, biosynthesis. The positive effect of *pykA* silencing can be explained from the fact that the absence of pyruvate kinase leads to more phosphoenolpyruvate to be converted to oxaloacetate by PEP carboxylase. More oxaloacetate can in return sustain a higher aspartate flux through AspC, which is crucial from threonine since, as already stated, threonine biosynthesis starts from aspartate. No further production of threonine was measured after more than 24 hours since Cas9 induction. This is expected since threonine is subject to a strong negative feedback regulation with key enzyme aspartokinase being inhibited and its expression repressed by threonine [4].

### Production of DMP from threonine

First, we proceed to test the conversion of threonine to DMP in *Escherichia coli*. We transformed *Escherichia coli* with a high-copy plasmid to express threonine dehydrogenase (which catalyzes the production of L-2-aminoacetoacetate from threonine), and aminoacetone oxidase from *Streptococcus oligofermentans*. Threonine de-hydrogenase is naturally present in *Escherichia coli* but is poorly expressed, which is why we decided to express it using the plasmid [16]. Cells were grown 24 hours in the self-inducing medium ZYM-5052 [16]. After 24 h the cells were collected via centrifugation, resuspended in 1/5 of the original volume in Tris HCl 20 mM pH 8 supplemented with 5 g/L threonine. Unfortunately, after 24 h no DMP production was detected. We speculated that the reason was the lack of threonine transporters at the time of biomass recovery after 24 h of growth or lack of activity of the transporters in buffer. For example, the H^+^-dependent TdcC system is active exclusively under anaerobic conditions in rich medium, while the major threonine importer, SstT, is dependent from Na^+^ which is not present in the reaction [17] [18]. For this reason, we decided to disrupt the cells after resuspension in the buffer and to perform biocatalysis using the cell extract. This led to positive results, with a DMP yield of around 20% (mol/mol). The are shown in Figure 4.

**Figure 4:**
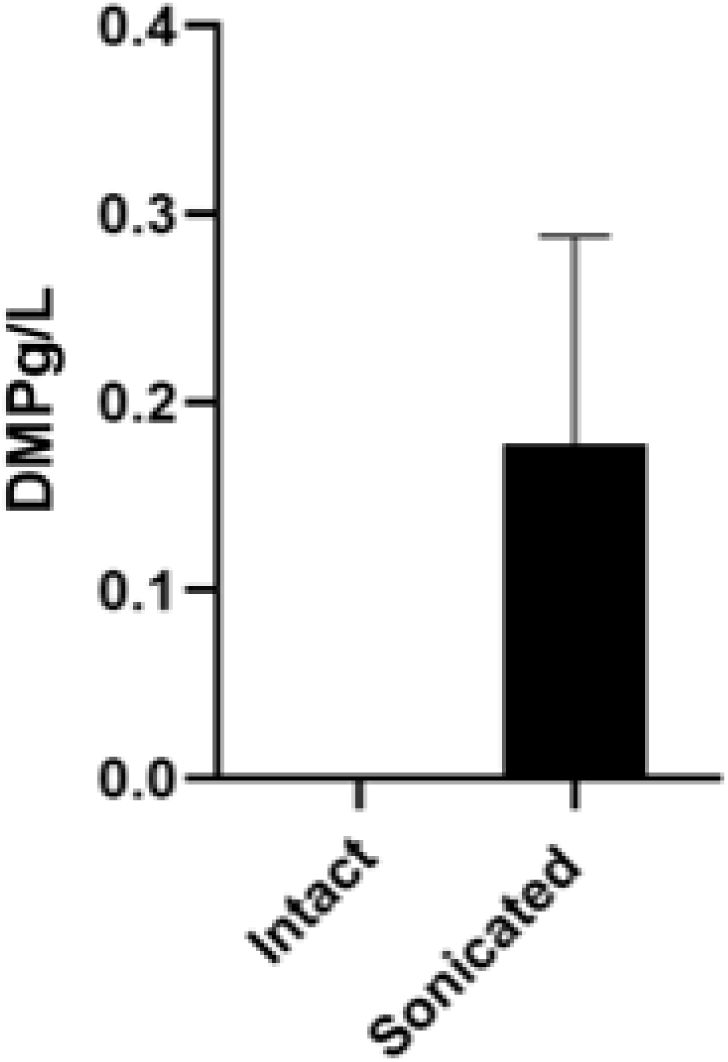
Testing the production of DMP from threonine in intact and sonicated biomass. In the first case there was no production, possibly due to lack of threonine importers at the moment of biomass recover.

**Figure 5:**
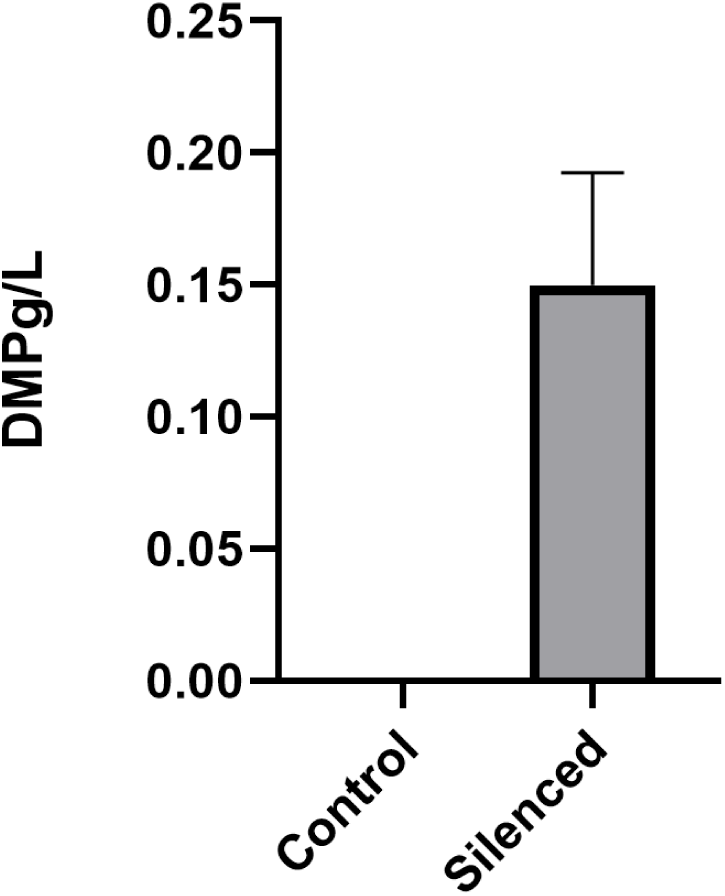
Production of DMP from glucose using the strain for increased threonine biosynthesis and the cell extract from the strain expressing threonine dehydrogenase and aminoacetone oxidase.

After we positively tested the activity of threonine dehydrogenase and aminoacetone oxidase, we proceeded to join the two strategies by producing DMP exploiting the increased threonine production flux. The experiment was performed in the same way as previously stated for CRISPi, but this time IPTG was also added for induction of threonine dehydrogenase and aminoacetone oxidase. After 24 hours, the supernatant was sampled for DMP quantification. As control, we used the strain with all the machinery for silencing without inducing Cas9. DMP was present in the surnatant of the induced cells, although the yield on glucose was low (0.3% mol/mol), which underlies the crucial role of increasing the threonine synthesis flux.

## Conclusions

Using Flux Balance Analysis, we rapidly identified phosphoserine phosphatase, phosphoserine deaminase and pyruvate kinase as genes of interest for increased threonine production. CRISPRi-mediated gene silencing confirmed these predictions, with a 5.6-fold increase of threonine production compared to the control. Therefore, we were able to identify a novel strategy for increased threonine flux solely by computational analysis of the genome-wide model of *Escherichia coli*, while other approaches all relied on either experimental data or from conclusion drawn from the literature on the subject (for example, see [2] [3] [4] [5] [6]). Subsequently, to show the potential of this new strategy we used it in conjunction with expression of threonine dehydrogenase and aminoacetone oxidase to produce the high value-added chemical 2,5-dimethylpyrazine straight from glucose. Although production is lower than other data reported in the literature [1], production of DMP showed the potential of the new strategy. This silencing strategy could be used in conjunction with known approaches to develop even better threonine and DMP producing *Escherichia coli* strains.

## Acknowledgements

This work is part of a public-private partnership with Anonima Materie Sintetiche e Affini SpA in the context of the Piano Nazionale Resistenza e Resilienza Mission 4 - Componente 1 - Riforma 4.1 Riforma dei Dottorati – Inv. 4.1 Borse PNRR patrimonio Culturale. Marco Vanoni also acknowledges the “ELIXIRxNextGenerationIT” (Code IR0000010)-CUP B53C22001800006 to Marco Vanoni. Funding was provided by the Euorpean Union’s project “Next Generation EU”.

## Notes

### Competing Interest Statement

The authors have declared no competing interest.

